# Constitutional activation of BMP4 and WNT signalling in hESC results in impaired mesendoderm differentiation

**DOI:** 10.1101/2020.06.04.133496

**Authors:** C. Markouli, E. Couvreu De Deckersberg, D. Dziedzicka, M. Regin, S. Franck, A. Keller, A. Gheldof, M. Geens, K. Sermon, C. Spits

## Abstract

We identified a human embryonic stem cell subline that fails to respond to the differentiation cues needed to obtain mesendoderm derivatives, differentiating into trophoblast-like cells instead. RNA-sequencing analysis showed that the subline has hyperactivation of the WNT and BMP4 signalling. Modulation of these pathways with small molecules confirmed them as the cause of the differentiation impairment. While activation of WNT and BMP4 in control cells resulted in a loss of mesendoderm differentiation and induction of trophoblast markers, inhibition of these pathways in the subline restored its ability to differentiate. Karyotyping and exome sequencing analysis did not identify any changes in the genome that could account for the pathway deregulation. These findings add to the increasing evidence that different responses of stem cell lines to differentiation protocols are based on genetic and epigenetic factors, inherent to the line or acquired during cell culture.

## Introduction

Human embryonic stem cells (hESCs) are a potent tool for the study of early development, in disease modelling and in regenerative medicine, as they can differentiate to any cell type of the human body. HESC differentiation protocols aim to mimic the fine orchestration of time dependent pathway modulation which is observed *in vivo*. Given this complexity, it is broadly acknowledged that individual cell lines may display a preference for differentiation to one germ layer over another (Kim et al., 2017; Osafune et al., 2008). This differentiation bias appears to be modulated by a wide range of factors, including genetic and epigenetic abnormalities (reviewed in (Keller et al., 2018)).

Prolonged culture affects the genetic integrity of hESC lines, resulting in point mutations (Merkle et al., 2017) and chromosomal abnormalities ranging from whole chromosome aneuploidies to smaller structural variants (Keller et al., 2018; Nguyen et al., 2013). Common aneuploidies are entire or partial gains of chromosomes 1, 12, 17, and X (Amps et al., 2011; Baker et al., 2007; Draper et al., 2004; Herszfeld et al., 2006; Maitra et al., 2005; Mitalipova et al., 2005; Yang et al., 2010) while a gain of 20q11.21 is found in more than 20% of stem cell lines worldwide (Amps et al., 2011; Avery et al., 2013; Nguyen et al., 2014). The selective advantage driving these abnormalities, and their potential impact on differentiation is poorly understood. As an exception, the gain of 20q11.21 leads to a *BCL2L1* dependent decrease in apoptotic sensitivity (Avery et al., 2013; Nguyen et al., 2014) and an impaired TGF-β-dependent neuroectodermal commitment (Markouli et al., 2019). Furthermore, hESC carrying an extra copy of chromosome 12 proliferate faster and display a limited differentiation capacity leading to the presence of undifferentiated cells in teratomas (Ben-David et al., 2014).

Epigenetic variance also plays an important role in differentiation bias (Adewumi et al., 2007; Bar and Benvenisty, 2019; Lagarkova et al., 2006). These changes vary from DNA methylation or histone modifications (Gifford et al., 2013; Xie et al., 2013), to chromatin remodelling (Dixon et al., 2015) and X-chromosome inactivation (Geens and Chuva De Sousa Lopes, 2017; Salomonis et al., 2016). For instance, mesoderm initiation requires low levels of H3K27me3 histone marks (Wang et al., 2017) while loss of H3K4me3 facilitates neuroectoderm differentiation (Bertero et al., 2015), and decreased global non-CG DNA methylation correlates with low endodermal differentiation capacity (Butcher et al., 2016). Additionally, expression levels of miR-371-3 and other non-coding RNAs have been linked to decreased neuroectoderm differentiation efficiency (Kim et al., 2011; Mo et al., 2015), and WNT3 levels are positively correlating with definitive endoderm commitment (Jiang et al., 2013a). Lastly, *RUNX1A* affects hematopoietic lineage formation (Ran et al., 2013) and β*FGF-1*, *RHOU* and *TYMP* are associated with low hepatic differentiation efficiency (Yanagihara et al., 2016).

All these findings point to differentiation bias as a complex phenomenon modulated by numerous possible genetic/epigenetic variations. In this study we investigate the mechanisms behind the loss of differentiation capacity of one of the hESC sublines in our laboratory. We identified a subline, here named VUB03_S2, with a strongly impaired differentiation capacity toward mesendodermal derivates, and developing a trophoblast-like expression profile. Here we show that this is due to constitutionally hyperactivated BMP4 and WNT signalling and confirm our findings by changing cell fate in control hESC when subjecting them to the same signalling cues during mesendoderm differentiation.

## Results

### VUB03_S2 shows impaired differentiation to all three lineages and differentiates to trophoblast-like cells under definitive endoderm differentiation conditions

VUB03_S2 is a genetically abnormal hESC sub-line in our lab, that has acquired a gain of 20q11.21. We first assessed the differentiation capacity of VUB03_S2 to neuroectoderm, as compared to its genetically normal counterpart VUB03_S1. VUB03_S2 differentiated poorly to neuroectoderm after a 4-day induction protocol using a dual TGF-β inhibition (Chambers et al., 2009; Chetty et al., 2013). We observed a 140-fold higher *PAX6* mRNA expression in VUB03_S1-derived cells and 0,65% PAX6^+^ cells in the case of VUB03_S2 (Figure 1A). These results are in line with our recently published work showing that hESC with a gain of 20q11.21 have a TGF-β-dependent impaired neuroectoderm commitment (Markouli et al., 2019).

**Figure 1.**
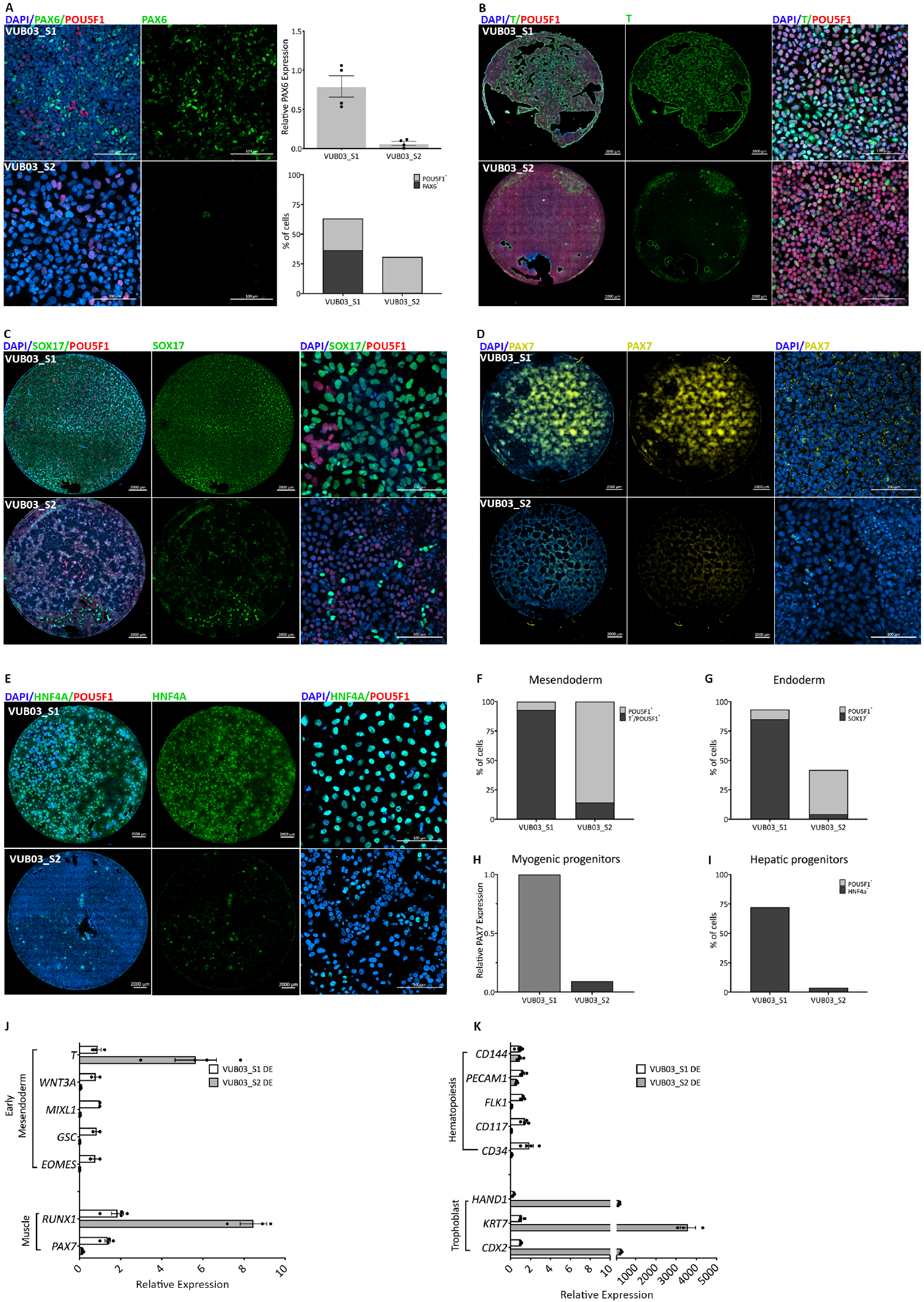
VUB03_S2 shows impaired commitment towards all three germ layers and upregulates trophoblast markers during definitive endoderm differentiation. A (left panels: immunostaining for PAX6 (green) and POU5F1 (red) in VUB03_S1 and S2 after 4 days of neuroectoderm differentiation. A (top right panel): Expression of the neuroectoderm marker *PAX6* relative to VUB03_S1 (n=4). Data are shown as mean ±SEM, each dot represents an independent differentiation experiment and the horizontal bars with asterisks represent statistical significance between samples (P<0.05, t-test). A (bottom right panel): Counts for PAX6+ and POU5F1+ cells in neuroectoderm derived from VUB03_S1 and VUB03_S2. B: Immunostaining for T (green) and POU5F1 (red) in both lines after 1-day mesendoderm induction. The left and middle panels represent full-well images. C: Immunostaining for SOX17 (green) and POU5F1 (red) in both lines after 3-day definitive endoderm differentiation. The left and middle panels represent full-well images. D: Immunostaining for PAX7 (yellow) in both lines after 12-day striated myogenic progenitor differentiation. E: Immunostaining for HNF4A (green) and POU5F1 (red) in both lines after 8-day hepatoblast differentiation. (F, G, I) Counts of positive cells for each immunostaining. (H) *PAX7* mRNA expression relative to VUB03_S1. Expression of (J) early mesendoderm and muscle, (K) hematopoietic lineage and trophoblast markers after 3-day definitive endoderm differentiation relative to VUB03_S1 (n=3 to 4). Data are shown as mean ±SEM and each dot represents an independent differentiation experiment. Scale bars represent 2000μm and 100μm.

Next, we assessed the differentiation to mesendoderm through a 24h induction that mimics the first steps of primitive streak formation, by WNT and TGF-β activation. We found that 93% of VUB03_S1 cells versus 14% of VUB03_S2 are T^+^ (a marker for early mesendoderm specification), while for both lines most cells are also POU5F1^+^ (Figure 1B and 1F). These results show that while VUB03_S1 can readily be induced into the mesendoderm lineage, VUB03_S2 mostly remains undifferentiated.

We then performed a 3-day differentiation to definitive endoderm through WNT and TGF-β activation for the first 24h and only TGF-β activation thereafter. As seen during the mesendoderm induction, VUB03_S1 efficiently differentiates to endoderm, with 85% of cells being SOX17^+^, while VUB03_S2 shows 4% of correctly differentiated cells. Eight percent of VUB03_S1 cells and 38% of VUB02_S2 were POU5F1^+^ (Figure 1C, G). This suggests that while the ability of VUB03_S2 to commit to mesendoderm is severely impaired, the cells are able to exit the pluripotent state, albeit less efficiently.

To assess whether the deficiency was unique to the initial events of the primitive streak formation or whether this effect was also present in the later stages, we carried out longer differentiation protocols for both lineages. We performed a 12-day induction to myogenic progenitors with myogenic medium supplemented with CHIR99021 (WNT signalling activator) for the first 48h and subsequently by FGF2 for the remainder of the differentiation. As expected, VUB03_S2 differentiates poorly to myogenic progenitors, with an 11-fold lower yield in skeletal muscle marker *PAX7* transcripts, but it does exit the pluripotent state (Figure 1D, H). Lastly, we performed an 8-day differentiation to hepatic progenitors to confirm whether the same deficiency prevails for endoderm derivates. The immunofluorescence results for HNF4A (Figure 1E) and cell counts (Figure 1I) support that here too VUB03_S2 fails to differentiate to the endodermal lineage.

Taken together, the results show that VUB03_S2 has a severe impairment in mesendodermal differentiation, independent of the time that cells are exposed to differentiation signals. Also, VUB03_S2 did not remain undifferentiated, as indicated by the low expression of *POU5F1* in the differentiation protocols beyond day 1. In order to investigate the alternative cell fate reached by VUB03_S2, we assessed a variety of markers for different fates after 3 days of definitive endoderm differentiation: early mesendoderm (*T, WNT3A, MIXL1, GSC, EOMES*), muscle (*RUNX1, PAX7*), hematopoietic lineage (*CD144, PECAM, FLK1, CD117, CD34*) and trophoblast (*HAND1, KRT7, CDX2*)(Figure 1J, K). Of these markers, only those specific to trophoblast showed a significant upregulation in VUB03_S2 (500 to 3500 times) with the protein levels of KRT7 also found to be elevated (Supplementary Figure 1). We therefore concluded that VUB03_S2 does not respond as expected to differentiation cues, and instead preferentially differentiates to trophoblast-like cells.

### VUB03_S2 shows a distinct transcriptomic profile with a deregulation of BMP4 and WNT signalling

In order to investigate the mechanisms behind the lack of ability to differentiate to mesendoderm derivates, we carried out a transcriptome analysis by RNA sequencing on VUB03_S2 and four control lines with proven ability to differentiate to all three germ layers (VUB01, VUB02, VUB03_S1 and VUB14, differentiation data for VUB01 and VUB02 published in Markouli et al., 2019; data for VUB14 not shown). We included two to five replicates per line, obtained from samples collected from independent cell cultures. Only coding genes with a count per million greater than one in at least two samples were included in our analysis. In unsupervised hierarchical clustering, all five replicates of VUB03_S2 cluster together and apart from the other four lines, including VUB03_S1 (Figure 2A). The principal component analysis plot of all expressed genes shows the same pattern, with the five VUB03_S2 replicates organizing in a common cluster that is distinct from the control lines (Figure 2B). This illustrates that the transcriptome of VUB03_S2 differs significantly from the other tested lines. Differential gene expression analysis with a cut-off value of |log_2_ Fold Change|>1 and false discovery rate q-value (FDR)<0.05 shows 623 up-regulated and 888 down-regulated genes in VU03_S2 versus control lines (Figure 2C). A list with the top-50 differentially expressed genes can be found in the supplementary table 1.

**Figure 2.**
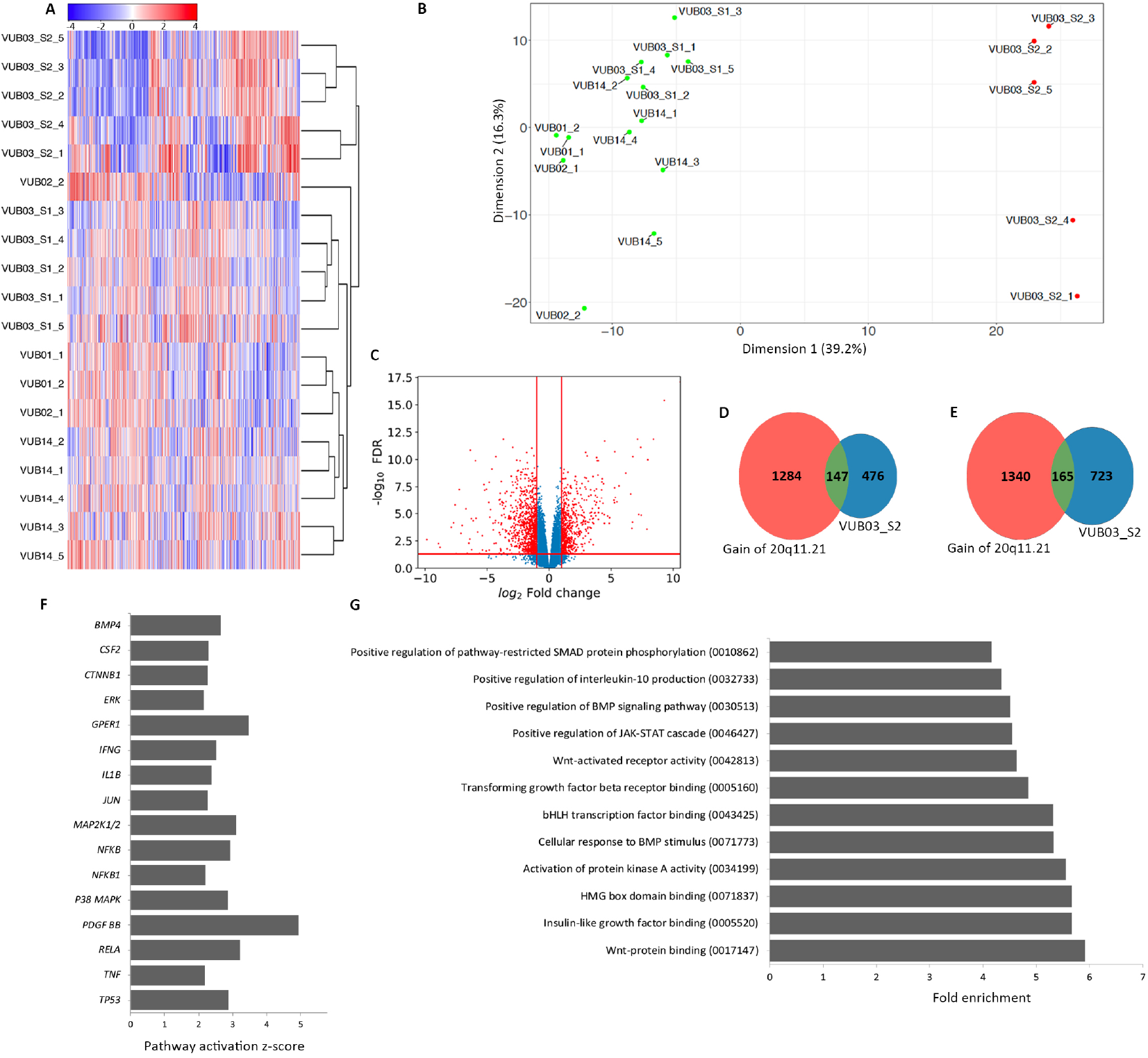
VUB03_S2 shows a unique transcriptomic signature characterized by WNT and BMP4 signalling deregulation. (A) Unsupervised Cluster Analysis and (B) Principal component analysis of component 1 versus component 2 of all coding genes with a count per million greater than one in at least two samples. (C) Volcano plot of the differential gene expression analysis of VU03_S2 versus all other hESC lines. The red lines show cut-off values of |log2 Fold Change|>1 and FDR<0.05. (D, E) Venn diagrams comparing the deregulated genes in VUB03_S2 with those deregulated in lines with a 20q11.21 gain. (D) Shows the upregulated genes and (E) the downregulated genes. (F) Ingenuity Pathway Analysis of all differentially expressed genes with |log_2_ Fold Change|>1 and FDR<0.05. The plot shows only upstream regulators of pathways with an activation score below −2 and above 2 with a p<0.05. (G) DAVID Enrichment Analysis of the 1000 top-deregulated genes, showing only gene-ontology terms related to intracellular signalling (full list in supplementary table 2). In parenthesis are the gene-ontology term numbers.

To ensure that these differences were not the result of the gain of 20q11.21 present in VUB03_S2, we compared the up- and down-regulated genes found in VUB03_S2 to those identified in other lines with a gain of 20q11.21 (Figure 2D and 2E respectively, data published in Markouli *et al.,* 2019). Thirty percent of the upregulated genes and 22% of the downregulated genes in VUB03_S2 are in common to those deregulated in lines with a gain of 20q11.21. This limited transcriptomic similarity between the two groups shows that VUB03_S2 displays a unique transcriptome that is only partially related to the chromosomal abnormality it carries.

We used Ingenuity Pathway Analysis (Krämer et al., 2014), using all genes with a |log_2_ Fold Change|>1 and FDR<0.05, to predict the activation state of upstream pathway regulators. We considered only pathways with p-value<0.05 and |z-score|>2, which are predicted by 10 or more regulators to be significantly activated or inhibited. (Figure 2F). We also carried out functional annotation enrichment analysis of the top-1000 deregulated genes using DAVID. This retrieved a list of 558 annotations, of which we selected the gene ontology terms with a fold-enrichment >4, and a p-value <0.05. The list of gene ontology terms referring to intracellular signalling is shown in Figure 2G, the full list of 49 terms can be found in the supplementary table 2. Based on the common predictions retrieved from both tools, we could conclude that VUB03_S2 shows differentially regulated pathways that are key to the control of differentiation such as BMP4 and WNT (beta-catenin).

### The transcriptomic profile of VUB03-S2 cannot be explained by additional chromosomal abnormalities or exomic point mutations

As mentioned above, VUB03_S2 carries a 4Mb gain of 20q11.21, with no further *de novo* gains or losses in other chromosomes. Supplementary table 3 contains the karyotypes of all lines used in the study. The gain of 20q11.21 in VUB03_S2 unlikely explains the phenotype since in our lab we have studied multiple hESC lines with smaller and larger gains of 20q11.21 and with higher copy numbers, none of which showed signs of poor mesendoderm commitment (Markouli et al., 2019). This suggests that VUB03_S2, although karyotypically similar to these lines, presents a unique differentiation bias unrelated to its karyotype.

In order to investigate if *de novo* point mutations could be responsible for the transcriptomic changes observed in VUB03_S2, we carried out whole exome sequencing on bulk DNA samples of VUB03_S1 and VUB03_S2. We first filtered the variants for de novo appearance in VUB03_S2 and by only considering those with a read depth >10. Recurrent sequencing artefacts were avoided by omitting changes that are found >100 times in the exome sequencing database of our sequencing facility. Finally, we only considered variants that were present in at least 30% of reads, to only include variants that were fixed in the population in at least a heterozygous state. This yielded 917 variants, of which 849 were in non-coding regions and 68 in protein coding sequences (supplementary table 4 shows an overview of the location of the variants). Fifty-two of the protein-coding variants were non-synonymous. The exact location and nature of these variants are listed in the supplementary table 5. We annotated the functions of the genes using the public databases NCBI-Gene and GeneCards, and found that none of the variants was located in a gene with a link to the WNT and BMP4 signalling.

Alternatively, the loss of a transcription factor binding site could potentially explain changes in gene-expression and pathway activation. As such we evaluated the possibility that a variant had negatively impacted transcription factor binding sites in *BMP4*, *CTNNB1* and their downstream effectors and targets (list obtained from the Ingenuity Pathway Analysis database, 198 and 535 genes for *BMP4* and *CTNNB1*). We found that 13 of these genes contained a *de novo* variant (supplementary table 6). We used cistrome (Vorontsov et al., 2018) to identify the transcription factors binding the regions containing the variants and PWMScan to establish if the variant abolished the transcription binding site (Ambrosini et al., 2018). None of the variants resulted in the destruction of transcription factor binding sites. Because of the use of exome sequencing data, we could only assess a limited number of transcription binding sites. Therefore, it cannot be excluded that the subline carries variants in regulatory regions not covered by our sequencing approach.

### Sustained intrinsic WNT and BMP4 activation impairs differentiation to definitive endoderm and drives the cells towards a trophoblast-like fate

The transcriptomic analysis predicted dysregulation of the WNT and BMP signalling, both are key regulators of differentiation and lineage commitment. A brief WNT activation is necessary to induce definitive endoderm differentiation, and the most common approach of inducing differentiation towards trophoblast-like cells is BMP4 treatment (reviewed in Gamage et al., 2016). Therefore, we aimed at validating the implication of these pathways in the misspecification observed in VUB03_S2. First, we mimicked the phenotype by subjecting control lines to the standard differentiation media, supplemented with recombinant BMP4 and the WNT activator CHIR99021 (experimental setup illustrated in Figure 3A). The identification of the optimal concentrations of factors was established using VUB03_S1, and their effect validated on two additional control lines.

**Figure 3.**
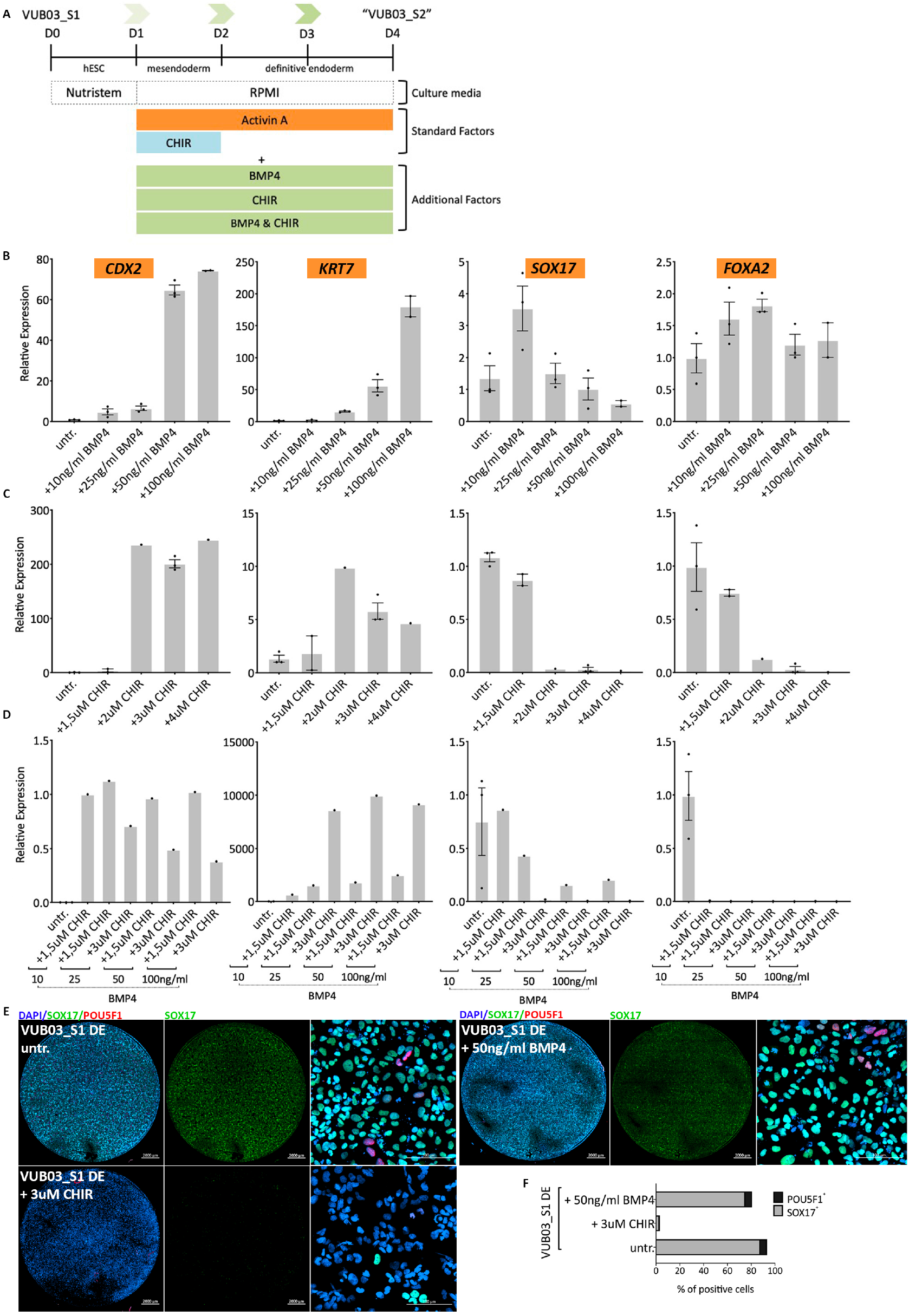
Impairing the definitive endoderm differentiation of VUB03_S1 through WNT and BMP4 pathway activation. (A) A schematic representation of the experimental design. WNT and BMP4 activators, in different concentrations, are added to the normal definitive endoderm differentiation medium. (B-D) mRNA expression of trophoblast markers *CDX2, KRT7* and definitive endoderm markers *SOX17* and *FOXA2* relative to VUB03_S1 after definitive endoderm differentiation. (B) After use of BMP4 only. (C) After use of CHIR99021 only. (D) Combination of both factors. (n=1 to 3) Data are shown as mean ±SEM, each dot represents an independent differentiation experiment. (E) Immunostaining for SOX17 (green) and POU5F1 (red) and (F) counts for SOX17 and POU5F1-positive cells in VUB03_S1 after 3 days of definitive endoderm differentiation untreated and after BMP4 and CHIR treatment. Scale bars represent 2000μm and 100μm.

First, we exposed the cells to exogenous BMP4 during definitive endoderm differentiation, in four different concentrations: 10ng/ml, 25ng/ml, 50ng/ml and 100ng/ml (Figure 3B). We evaluated mRNA levels for *CDX2* and *KRT7* to determine if the cells were becoming trophoblast-like and *SOX17, FOXA2* to evaluate definitive endoderm. While the addition of BMP4 did not affect the expression levels of the specific definitive endoderm markers, *CDX2* and *KRT7* were induced at high concentrations (63 and 75-fold increase as compared to the untreated cells). These findings suggest that BMP4 activation is necessary for the cells to acquire a trophoblast-like expression profile while having little effect on the expression of DE markers. This is further illustrated in Figure 3E and 3F where the SOX17 protein levels between the untreated and BMP4-treated VUB03_S1 show minimal differences.

We then tested the effect of WNT activation using different concentrations of CHIR99021 (1.5, 2, 3 and 4μM, Figure 3C). While the lowest concentration (1.5 μM) of CHIR did not show a noticeable effect compared to the untreated line, the three higher concentrations (2, 3 and 4μM) resulted in a significant induction of *CDX2* expression (200-fold increase), a modest increase in the levels of *KRT7* (7-fold increase) and strong decrease in *SOX17* and *FOXA2* (80-300-fold decrease). Nevertheless, although *KRT7* was up-regulated in comparison to the untreated cells, the induction was lower than what we observed after the addition of BMP4. The effect on definitive endoderm induction was also demonstrated at the protein level in Figure 3E and 3F, with a strong decrease of SOX17^+^ cells in the CHIR-treated condition compared to the untreated.

When we combined both factors (Figure 3D), we found that the combination of 3μM of CHIR and any amount of BMP4 between 25 and 100ng/mL resulted in a loss of endoderm specification and a differentiation into trophoblast-like cells. This suggests that WNT activation has the strongest effect on the impairment of differentiation to definitive endoderm while BMP4 is mostly responsible for the *KRT7* up-regulation, revealing a synergistic effect of the two pathways to explain the phenotype of VUB03_S2. We confirmed these results on two additional karyotypically normal lines (VUB02 and VUB19), treating them with 3μM CHIR and 50ng/mL BMP4 (Supplementary Figure 2 and 3).

Finally, to validate our findings, we aimed to rescue the definitive endoderm differentiation capacity of VUB03_S2 by exposure to noggin (NG, BMP4 pathway inhibitor) and XAV-939 (WNT pathway inhibitor). The experimental design is shown in Figure 4A. At the concentrations tested, Noggin did not inhibit the induction of *CDX2* and *KRT7* but did increase the expression of *FOXA2* (9 to 19-fold) while the effect on *SOX17* was modest (1.5-2-fold increase). On the other hand, treatment with XAV-939 resulted in 3 to 19-fold decrease in both trophoblast markers and a significant improvement in the definitive endoderm differentiation (Figure 4C) while combination of both factors showed a similar outcome (Figure 4D). The effect of each compound on DE induction can be also seen in the immunostaining images for SOX17, in VUB03_S2, Noggin and XAV-treated cells (Figure 4E). Cell counts for SOX17^+^ and POU5F1^+^ cells (Figure 4F) suggest that WNT inhibition of VUB03_S2 with XAV-939 rescues its DE deficiency and restores SOX17 levels comparable to VUB03_S1 untreated cells (Figure 3F).

**Figure 4.**
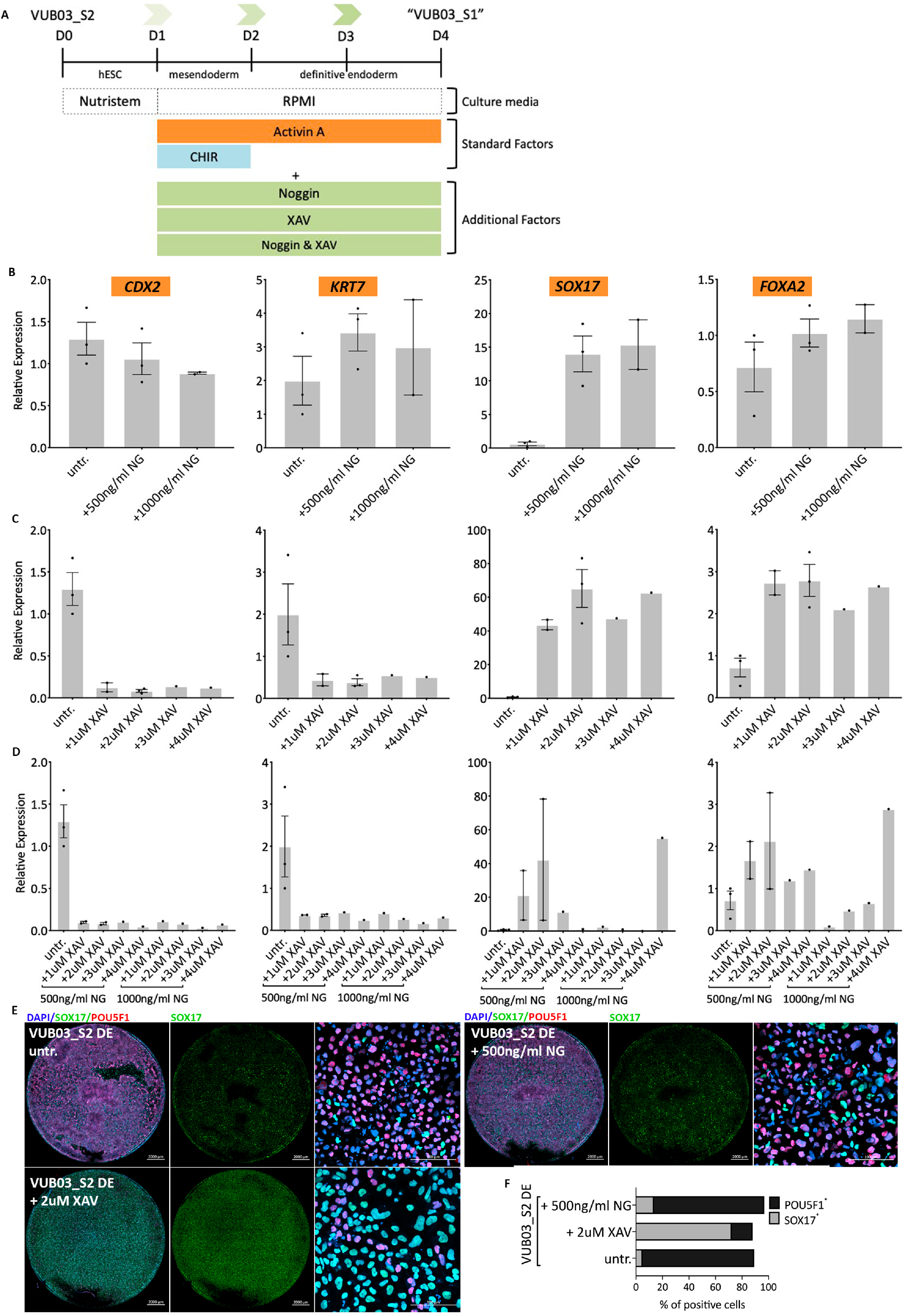
Restoration of the definitive endoderm differentiation capacity of VUB03_S2 through WNT and BMP4 pathway inhibition. (A) A schematic representation of the experimental design. WNT and BMP4 inhibitors, in different concentrations, are added to the normal definitive endoderm differentiation (B-D) mRNA expression of trophoblast markers *CDX2, KRT7* and definitive endoderm markers *SOX17* and *FOXA2* relative to VUB03_S2 untreated cells after definitive endoderm differentiation. (B) After use of Noggin only. (C) After use of XAV-939 only. (D) Combination of both factors (n=1 to 3). Data are shown as mean ±SEM, each dot represents an independent differentiation experiment. (E) Immunostaining for SOX17 (green) and POU5F1 (red) and (F) counts for SOX17 and POU5F1-positive cells in VUB03_S2 untreated / Noggin and XAV-treated cells after 3 days of definitive endoderm differentiation. Scale bars represent 2000μm and 100μm.

## Discussion

The aim of our study was to elucidate the cause of the impaired differentiation capacity of VUB03_S2. We previously showed that the gain of 20q11.21 this subline carries causes the failure to differentiate to neuroectoderm (Markouli et al., 2019), but this could not explain its reduced mesendoderm differentiation capacity. Moreover, this subline is able to exit pluripotency, and to preferably yields trophoblast-like cells. In this study we demonstrate that intrinsically hyperactivated WNT and BMP4 signalling are responsible for this atypical response to mesendoderm differentiation cues. Moreover, we were able to reverse this response by the use of WNT and BMP4 inhibitors.

To the best of our knowledge, this is the first report on a human pluripotent stem cell (hPSC) line that is not only showing differentiation impairment, but also specification to the wrong lineage when submitted to well-established differentiation cues. Several groups have reported that the expression levels of specific genes may modulate the propensity of a line to a specific lineage, but in none of these cases the lines committed to a completely different lineage (Jiang et al., 2013a; Kim et al., 2011; Mo et al., 2015; Ran et al., 2013; Yanagihara et al., 2016). Here, we find that WNT and BMP4 activation, along with TGF-β activation by Activin A results in trophoblast-like cell formation. This finding is not entirely unexpected, as it is known that *CDX2* is a downstream target of WNT signaling (Sherwood et al., 2011; Vaninetti et al., 2009). Furthermore, BMP4 regulates the specification towards the trophoblast state (Jain et al., 2017) and is required for induction of *KRT7* expression in extraembryonic mesoderm and trophoblast cells (Bernardo et al., 2011).

While we could demonstrate the constitutional activation of these pathways resulting in the abnormal response to the differentiation cues, we could not identify its genomic cause. VUB03_S2 did not carry additional *de novo* chromosomal abnormalities or point mutations that could explain its transcriptomic profile, suggesting that the cause may either be found in the genomic regions that are not covered by the exome sequencing or potentially in epigenetic modifications. A number of studies have established a link between epigenomic changes and differentiation bias (Bock et al., 2011; Wang et al., 2017), although the mechanistic link remains to be unraveled.

Many current differentiation protocols suffer from reproducibility issues between laboratories and lines. While this is commonly thought to be a result of suboptimal protocols, it is becoming increasingly clear that individual lines are more likely at the orogin. Our work here shows in detail how hESC can acquire pervasive deregulation of intrinsic differentiation signals, thereby losing their ability to respond adequately to well established differentiation triggers. Furthermore, while a growing body of work has begun to demonstrate that failure to differentiate to certain lineages can be explained by commonly recurring mutations, often such outcomes may have no clear driver, as seen here, which may lead to the false conclusions on differentiation protocol reproducibility and general experimental variability. As hPSC are increasingly moving towards clinical trials, line choice will become increasingly important, and a comprehensive evaluation of differentiation biases will be vital to their long-term success.

## Experimental Procedures

### hESC lines and culture

HESC were derived and characterized as previously described (Mateizel et al., 2006, 2009). All lines are registered with the EU hPSC registry (https://hpscreg.eu/). hESC were cultured on dishes coated with 10μg/ml laminin-521 (Biolamina) in NutriStem hESC XF medium (NS medium; Biological Industries) with 100 U/ml penicillin/streptomycin (Thermo Fisher Scientific) and passaged as single cells in a 1:10 to 1:100 ration using TrypLE Express (Thermo Fisher Scientific) when 70-80% confluent.

### Differentiation to neuroectoderm

The protocol was adapted from (Chetty et al., 2013) and (Chambers et al., 2009). Differentiation was carried out on laminin-521 and initiated when the hESC culture was 90% confluent. The neurectoderm differentiation medium consisted of KnockOut D-MEM (Thermo Fisher Scientific), 10% KnockOut Serum Replacement (Thermo Fisher Scientific), 500ng/ml Recombinant Human Noggin Protein (R&D Systems) and 10μM SB431542 (Tocris) and was refreshed daily.

### Differentiation to mesendoderm and definitive endoderm

Endoderm differentiation was carried out using a protocol based on (Sui et al., 2012), on laminin-521. Differentiation was started when the cells were 80-90% confluent with medium containing Roswell Park Memorial Institute (RPMI) 1640 supplemented with GlutaMAX (Thermo Fisher Scientific), 0.5% B27 supplement (Thermo Fisher Scientific), 100 ng/ml Recombinant Human/Mouse/Rat Activin A (R&D Systems) and 3μM CHIR99021 (Stemgent). One day later the medium was changed to differentiation medium without CHIR99021 and the cells were cultured for two more days. For mesoderm induction, the same protocol was used but for only one day. To induce WNT and BMP4 activation we additionally supplemented the above protocol with CHIR99021 and/or Recombinant Human BMP4 Protein (R&D Systems) (figure 3A). To inhibit these pathways we used Recombinant Human Noggin Protein for BMP4 inhibition and/or XAV-939 for WNT inhibition (StemCell Technologies).

### Differentiation to hepatoblasts

Cells were plated in a ratio of 35.000cells/cm^2^ a day prior to differentiation. The next day the medium was changed to Roswell Park Memorial Institute (RPMI) 1640 medium supplemented by GlutaMAX (Thermo Fisher Scientific), 100 ng/ml Recombinant Human/Mouse/Rat Activin A and 3μM CHIR99021. One day later the medium was changed to differentiation medium without CHIR99021. From day 2 until day 8 the cells were cultured in Hepatoblast medium containing KnockOut D-MEM, 20% KnockOut Serum Replacement, 0.5X GlutaMAX, 1X 1% MEM Non-Essential Amino Acids (Thermo Fischer Scientific), 0.1mM β-Mercaptoethanol (Sigma Aldrich), and 1% DMSO (Sigma Aldrich). The medium was refreshed every day.

### Differentiation to Myogenic progenitors

Skeletal muscle differentiation was performed according to the protocol described in (Van Der Wal et al., 2017), with few adjustments. Briefly, a total of 50.000 cells/cm^2^ were plated on a laminin-521 coated dishes. The next day, differentiation was induced by the use of 10 µM CHIR99021 in myogenic differentiation medium consisting of DMEM-F12, 1x ITS-X, 100 U/ml Pen/Strep and 1x Glutamine (all from Thermo Fischer Scientific) for 2 days. Subsequently, the CHIR99021 was replaced by 20 ng/ml FGF2 (Prepotech) for the following 10 days. Medium was refreshed daily.

### Quantitative real-time PCR

RNA extraction was performed with the QIAGEN mini or micro RNAeasy kit, cDNA conversion was performed with the GE-healthcare cDNA synthesis kit, following the manufacturers’ instructions. qRT-PCR was carried out on a ViiA 7 thermocycler (Thermo Fisher Scientific) and using standard protocol as provided by the manufacturer. Details on the probes, assays and primers are listed in the supplementary table 7.

### Immunostaining

Immunostaining for neuroectoderm, mesoderm, definitive endoderm and hepatoblasts was carried out on cells fixed and permeabilized with 4% paraformaldehyde (PFA) and 100% Methanol (Sigma-Aldrich), and blocked with 10% Fetal Bovine Serum (Thermo Fischer Scientific). For the myogenic progenitor differentiation cells were fixed with (PFA) for 10 min at room temperature and permeabilized using 0.3% Triton-X (Sigma-Aldrich) and blocked with 3% BSA, 0.1% Tween in PBS. The list with antibodies and manufacturers can be found in the supplementary table 8. Primary antibodies were incubated overnight at 4^0^C, secondary antibodies were incubated for 2-3h at room temperature. Nuclear staining was performed with Hoechst 33342 (Thermo Fischer Scientific). Imaging was performed on a LSM800 confocal microscope (Carl Zeiss), and cell counts were done using the Zen 2 (blue edition) imaging software.

### RNA sequencing

150ng of RNA was used to perform an Illumina sequencing library preparation using the QuantSeq 3' mRNA-Seq Library Prep Kits (Lexogen) according to manufacturer's protocol. During library preparation 17 PCR cycles were used. Libraries were quantified by qPCR, according to Illumina's protocol 'Sequencing Library qPCR Quantification protocol guide', version February 2011. A High sensitivity DNA chip (Agilent Technologies) was used to control the library's size distribution and quality. Sequencing was performed on a high throughput Illumina NextSeq 500 flow cell generating 75 bp single reads. All data has been deposited on Gene Expression Omnibus repository with accession number GSE134454. The reads were mapped against the Genome Reference Consortium Human Build 38 patch release 10 (GRCh38.p10) (Zerbino et al., 2018) using STAR (version 2.5.3)(Dobin et al., 2013). RNA-Seq by Expectation Maximization (RSEM)(Li and Dewey, 2011) software (version 1.3.0) was used to produce the count table. Of the 63967 Ensembl’s genes, only the somatic and X chromosomes’ coding genes were considered (19802 genes in total).

### RNA-seq analysis

The RNA-seq analysis was performed using the R software (version 3.6.1) with the edgeR (Robinson et al., 2010) and DESeq2 (Love et al., 2014) libraries. Only genes with a count per million (cpm) greater than 1 in at least two samples were considered. The raw counts were normalized using the trimmed mean of M values (Robinson and Oshlack, 2010) (TMM) algorithm. The normalized counts were then transformed in a log_2_ fold-change (log_2_FC) table with their associated statistics, p-value and false discovery rate (FDR). Genes with a |log_2_FC| > 1 and a FDR < 0.05 were considered as significantly differentially expressed. A |log_2_FC| > 1 means at least two times more or two times less transcript in the mutant group in comparison to the wild-type group.

The data was represented using a heatmap with an unsupervised hierarchical clustering and a principal component analysis using all expressed coding genes were included. Ingenuity Pathway Analysis (Krämer et al., 2014) (QIAGEN Inc.) was used for the pathway analysis based on the differential gene expression between groups. The pathway enrichment analysis was done using DAVID 6.8 (Huang et al., 2009a, 2009b). The top-1000 genes with the highest log_2_FC in absolute value were selected. In the different categories, only the GO terms were taken for account, only terms with p-value<0.05 were considered relevant.

### Array comparative genomic hybridization (aCGH)

Oligonucleotide aCGH was carried out based on the protocol provided by the manufacturer (Agilent Technologies). A total of 400ng of DNA was labeled with Cy3 while the reference DNA (Promega) was labelled with Cy5. The samples are hybridized on the microarray slide (4×44K Human Genome CGH Microarray, Agilent Technologies). The slides were scanned using an Agilent dual laser DNA microarray scanner G2566AA. Only arrays with a SD. ≤0.20, signal intensity >50, background noise <5 and a derivative log-ratio <0.2 were taken into account. Cutoff values were set at three consecutive probes with an average log_2_ ratio over 0.3 for gains and of −0.45 for loss.

### Exome sequencing

DNA was isolated following the manufacturers’ instructions (QIAGEN, Dneasy blood & tissue kit). DNA library preparation was performed with the Kapa HyperPrep kit (Roche, Basel, Switzerland), followed by exome target capturing with the SeqCap EZ exome probes V3 kit (Roche, Basel, Switzerland). Cluster generation was carried out on a cBOT system (Illumina, San Diego, CA) and samples were subsequently sequenced (paired end, 2×250bp) on a HiSeq1500 machine (Illumina, San Diego, CA). Bio-informatic analysis of the data was done with an in-house developed pipeline based on Picard Tools (Broad Institute, Cambridge, MA), the BWA aligner (Li and Durbin, 2010), GATK (Broad Institute, Cambridge, MA) and Alamut Batch (Interactive BioSoftware, Rouen, France). For the variant filtering, we used the software package Highlander (https://sites.uclouvain.be/highlander/) and used the BAM files as input.

## Acknowledgments

The authors acknowledge their colleagues from the human embryonic stem cell lab for the derivation and culture of the lines. This work was supported by the Fonds for Scientific Research in Flanders -1505814N- (Fonds Wetenschappelijk Onderzoek – Vlaanderen (FWO)) and the Methusalem Grant to Karen Sermon, of the Research Council of the VUB. C.M is doctoral fellow supported by the Instituut voor Innovatie door Wetenschap en Technologie (IWT). A.K., E.C.D.D, M.R and D.D are doctoral fellows at the FWO.

## Author contributions

C.M. carried out all the experiments unless stated otherwise and co-wrote the manuscript. E.C.D.D. did the bioinformatics analysis. D.D. provided the control hESC samples for RNA sequencing. M.R. performed the microscopy and cell counting. S.F. performed the myogenic progenitor differentiation. A.K. assisted with hepatoblast differentiation. A.G. performed the exome analysis. K.S. edited the manuscript and provided funding. M.G. provided scientific advice and proofread the manuscript. C.S. co-wrote the manuscript, funded, designed and supervised the experimental work.

## References

Adewumi, O., Aflatoonian, B., Ahrlund-Richter, L., Amit, M., Andrews, P.W., Beighton, G., Bello, P.A., Benvenisty, N., Berry, L.S., Bevan, S., et al. (2007). Characterization of human embryonic stem cell lines by the International Stem Cell Initiative. Nat. Biotechnol. 25, 803–816.

Ambrosini, G., Groux, R., and Bucher, P. (2018). PWMScan: a fast tool for scanning entire genomes with a position-specific weight matrix. Bioinformatics 34, 2483–2484.

Amps, K., Andrews, P.W., Anyfantis, G., Armstrong, L., Avery, S., Baharvand, H., Baker, J., Baker, D., Munoz, M.B., Beil, S., et al. (2011). Screening ethnically diverse human embryonic stem cells identifies a chromosome 20 minimal amplicon conferring growth advantage. Nat. Biotechnol. 29, 1132–1144.

Avery, S., Hirst, A.J., Baker, D., Lim, C.Y., Alagaratnam, S., Skotheim, R.I., Lothe, R. a, Pera, M.F., Colman, A., Robson, P., et al. (2013). BCL-XL Mediates the Strong Selective Advantage of a 20q11.21 Amplification Commonly Found in Human Embryonic Stem Cell Cultures. Stem Cell Reports 1, 379–386.

Baker, D.E.C., Harrison, N.J., Maltby, E., Smith, K., Moore, H.D., Shaw, P.J., Heath, P.R., Holden, H., and Andrews, P.W. (2007). Adaptation to culture of human embryonic stem cells and oncogenesis in vivo. Nat. Biotechnol. 25, 207–215.

Bar, S., and Benvenisty, N. (2019). Epigenetic aberrations in human pluripotent stem cells. EMBO J. 38, e101033.

Ben-David, U., Arad, G., Weissbein, U., Mandefro, B., Maimon, A., Golan-Lev, T., Narwani, K., Clark, A.T., Andrews, P.W., Benvenisty, N., et al. (2014). Aneuploidy induces profound changes in gene expression, proliferation and tumorigenicity of human pluripotent stem cells. Nat. Commun. 5, 4825.

Bernardo, A.S., Faial, T., Gardner, L., Niakan, K.K., Ortmann, D., Senner, C.E., Callery, E.M., Trotter, M.W., Hemberger, M., Smith, J.C., et al. (2011). BRACHYURY and CDX2 Mediate BMP-Induced Differentiation of Human and Mouse Pluripotent Stem Cells into Embryonic and Extraembryonic Lineages. Cell Stem Cell 9, 144–155.

Bertero, A., Madrigal, P., Galli, A., Hubner, N.C., Moreno, I., Burks, D., Brown, S., Pedersen, R.A., Gaffney, D., Mendjan, S., et al. (2015). Activin/Nodal signaling and NANOG orchestrate human embryonic stem cell fate decisions by controlling the H3K4me3 chromatin mark. Genes Dev. 29, 702–717.

Bock, C., Kiskinis, E., Verstappen, G., Gu, H., Boulting, G., Smith, Z.D., Ziller, M., Croft, G.F., Amoroso, M.W., Oakley, D.H., et al. (2011). Reference Maps of Human ES and iPS Cell Variation Enable High-Throughput Characterization of Pluripotent Cell Lines. Cell 144, 439–452.

Butcher, L.M., Ito, M., Brimpari, M., Morris, T.J., Soares, F.A.C., Ährlund-Richter, L., Carey, N., Vallier, L., Ferguson-Smith, A.C., and Beck, S. (2016). Non-CG DNA methylation is a biomarker for assessing endodermal differentiation capacity in pluripotent stem cells. Nat. Commun. 7, 10458.

Chambers, S.M., Fasano, C.A., Papapetrou, E.P., Tomishima, M., Sadelain, M., and Studer, L. (2009). Highly efficient neural conversion of human ES and iPS cells by dual inhibition of SMAD signaling. Nat. Biotechnol. 27, 275–280.

Chetty, S., Pagliuca, F.W., Honore, C., Kweudjeu, A., Rezania, A., and Melton, D. a (2013). A simple tool to improve pluripotent stem cell differentiation. Nat. Methods 10, 553–556.

Dixon, J.R., Jung, I., Selvaraj, S., Shen, Y., Antosiewicz-Bourget, J.E., Lee, A.Y., Ye, Z., Kim, A., Rajagopal, N., Xie, W., et al. (2015). Chromatin architecture reorganization during stem cell differentiation. Nature 518, 331–336.

Dobin, A., Davis, C.A., Schlesinger, F., Drenkow, J., Zaleski, C., Jha, S., Batut, P., Chaisson, M., and Gingeras, T.R. (2013). STAR: ultrafast universal RNA-seq aligner. Bioinformatics 29, 15–21.

Draper, J.S., Smith, K., Gokhale, P., Moore, H.D., Maltby, E., Johnson, J., Meisner, L., Zwaka, T.P., Thomson, J. a, and Andrews, P.W. (2004). Recurrent gain of chromosomes 17q and 12 in cultured human embryonic stem cells. Nat. Biotechnol. 22, 53–54.

Gamage, T.K., Chamley, L.W., and James, J.L. (2016). Stem cell insights into human trophoblast lineage differentiation. Hum. Reprod. Update 23, 77–103.

Geens, M., and Chuva De Sousa Lopes, S.M. (2017). X chromosome inactivation in human pluripotent stem cells as a model for human development: back to the drawing board? Hum. Reprod. Update 23, 520–532.

Gifford, C. a., Ziller, M.J., Gu, H., Trapnell, C., Donaghey, J., Tsankov, A., Shalek, A.K., Kelley, D.R., Shishkin, A. a., Issner, R., et al. (2013). Transcriptional and epigenetic dynamics during specification of human embryonic stem cells. Cell 153, 1149–1163.

Herszfeld, D., Wolvetang, E., Langton-Bunker, E., Chung, T.-L., Filipczyk, A. a, Houssami, S., Jamshidi, P., Koh, K., Laslett, A.L., Michalska, A., et al. (2006). CD30 is a survival factor and a biomarker for transformed human pluripotent stem cells. Nat. Biotechnol. 24, 351–357.

Huang, D.W., Sherman, B.T., and Lempicki, R.A. (2009a). Bioinformatics enrichment tools: paths toward the comprehensive functional analysis of large gene lists. Nucleic Acids Res. 37, 1–13.

Huang, D.W., Sherman, B.T., and Lempicki, R.A. (2009b). Systematic and integrative analysis of large gene lists using DAVID bioinformatics resources. Nat. Protoc. 4, 44–57.

Jain, A., Ezashi, T., Roberts, R.M., and Tuteja, G. (2017). Deciphering transcriptional regulation in human embryonic stem cells specified towards a trophoblast fate. Sci. Rep. 7.

Jiang, W., Zhang, D., Bursac, N., and Zhang, Y. (2013a). WNT3 Is a Biomarker Capable of Predicting the Definitive Endoderm Differentiation Potential of hESCs. Stem Cell Reports 1, 46–52.

Jiang, W., Wang, J., and Zhang, Y. (2013b). Histone H3K27me3 demethylases KDM6A and KDM6B modulate definitive endodermdifferentiation from human ESCs by regulating WNT signaling pathway. Cell Res. 23, 122.

Keller, A., Dziedzicka, D., Zambelli, F., Markouli, C., Sermon, K., Spits, C., and Geens, M. (2018). Genetic and epigenetic factors which modulate differentiation propensity in human pluripotent stem cells. Hum. Reprod. Update 24, 162–175.

Kim, H., Lee, G., Ganat, Y., Papapetrou, E.P., Lipchina, I., Socci, N.D., Sadelain, M., and Studer, L. (2011). miR-371-3 expression predicts neural differentiation propensity in human pluripotent stem cells. Cell Stem Cell 8, 695–706.

Kim, S.K., Kim, J., Kim, S., Kim, B., Gil, J., Kim, S., and Kim, J. (2017). Comparative analysis of the developmental competence of three human embryonic stem cell lines in vitro Comparative Analysis of the Developmental Competence of Three Human Embryonic Stem Cell Lines in Vitro. 48–56.

Krämer, A., Green, J., Pollard, J., and Tugendreich, S. (2014). Causal analysis approaches in Ingenuity Pathway Analysis. Bioinformatics 30, 523–530.

Lagarkova, M.A., Volchkov, P.Y., Lyakisheva, A. V., Philonenko, E.S., and Kiselev, S.L. (2006). Diverse Epigenetic Profile of Novel Human Embryonic Stem Cell Lines. Cell Cycle 5, 416–420.

Li, B., and Dewey, C.N. (2011). RSEM: accurate transcript quantification from RNA-Seq data with or without a reference genome. BMC Bioinformatics 12, 323.

Li, H., and Durbin, R. (2010). Fast and accurate long-read alignment with Burrows-Wheeler transform. Bioinformatics 26, 589–595.

Love, M.I., Huber, W., and Anders, S. (2014). Moderated estimation of fold change and dispersion for RNA-seq data with DESeq2. Genome Biol. 15, 550.

Maitra, A., Arking, D.E., Shivapurkar, N., Ikeda, M., Stastny, V., Kassauei, K., Sui, G., Cutler, D.J., Liu, Y., Brimble, S.N., et al. (2005). Genomic alterations in cultured human embryonic stem cells. Nat Genet 37, 1099–1103.

Markouli, C., De Deckersberg, E.C., Regin, M., Nguyen, H.T., Zambelli, F., Keller, A., Dziedzicka, D., De Kock, J., Tilleman, L., Van Nieuwerburgh, F., et al. (2019). Gain of 20q11.21 in Human Pluripotent Stem Cells Impairs TGF-β-Dependent Neuroectodermal Commitment. Stem Cell Reports 0.

Mateizel, I., De Temmerman, N., Ullmann, U., Cauffman, G., Sermon, K., Van de Velde, H., De Rycke, M., Degreef, E., Devroey, P., Liebaers, I., et al. (2006). Derivation of human embryonic stem cell lines from embryos obtained after IVF and after PGD for monogenic disorders. Hum. Reprod. 21, 503–511.

Mateizel, I., Spits, C., Verloes, a, Mertzanidou, a, Liebaers, I., and Sermon, K. (2009). Characterization of CD30 expression in human embryonic stem cell lines cultured in serum-free media and passaged mechanically. Hum. Reprod. 24, 2477–2489.

Merkle, F.T., Ghosh, S., Kamitaki, N., Mitchell, J., Avior, Y., Mello, C., Kashin, S., Mekhoubad, S., Ilic, D., Charlton, M., et al. (2017). Human pluripotent stem cells recurrently acquire and expand dominant negative P53 mutations. Nature 545, 229–233.

Mitalipova, M.M., Rao, R.R., Hoyer, D.M., Johnson, J.A., Meisner, L.F., Jones, K.L., Dalton, S., and Stice, S.L. (2005). Preserving the genetic integrity of human embryonic stem cells. Nat. Biotechnol. 23, 19–20.

Mo, C.-F., Wu, F.-C., Tai, K.-Y., Chang, W.-C., Chang, K.-W., Kuo, H.-C., Ho, H.-N., Chen, H.-F., and Lin, S.-P. (2015). Loss of non-coding RNA expression from the DLK1-DIO3 imprinted locus correlates with reduced neural differentiation potential in human embryonic stem cell lines. Stem Cell Res. Ther. 6, 1.

Nguyen, H.T., Geens, M., and Spits, C. (2013). Genetic and epigenetic instability in human pluripotent stem cells. Hum. Reprod. Update 19, 187–205.

Nguyen, H.T., Geens, M., Mertzanidou, a, Jacobs, K., Heirman, C., Breckpot, K., and Spits, C. (2014). Gain of 20q11.21 in human embryonic stem cells improves cell survival by increased expression of Bcl-xL. Mol. Hum. Reprod. 20, 168–177.

Osafune, K., Caron, L., Borowiak, M., Martinez, R.J., Fitz-Gerald, C.S., Sato, Y., Cowan, C. a, Chien, K.R., and Melton, D. a (2008). Marked differences in differentiation propensity among human embryonic stem cell lines. Nat. Biotechnol. 26, 313–315.

Ran, D., Shia, W.-J., Lo, M.-C., Fan, J.-B., Knorr, D. a, Ferrell, P.I., Ye, Z., Yan, M., Cheng, L., Kaufman, D.S., et al. (2013). RUNX1a enhances hematopoietic lineage commitment from human embryonic stem cells and inducible pluripotent stem cells. Blood 121, 2882–2890.

Robinson, M.D., and Oshlack, A. (2010). A scaling normalization method for differential expression analysis of RNA-seq data. Genome Biol. 11, R25.

Robinson, M.D., McCarthy, D.J., and Smyth, G.K. (2010). edgeR: a Bioconductor package for differential expression analysis of digital gene expression data. Bioinformatics 26, 139–140.

Salomonis, N., Dexheimer, P.J., Omberg, L., Schroll, R., Bush, S., Huo, J., Schriml, L., Ho Sui, S., Keddache, M., Mayhew, C., et al. (2016). Integrated Genomic Analysis of Diverse Induced Pluripotent Stem Cells from the Progenitor Cell Biology Consortium. Stem Cell Reports 7, 110–125.

Sherwood, R.I., Maehr, R., Mazzoni, E.O., and Melton, D.A. (2011). Wnt Signaling Specifies and Patterns Intestinal Endoderm.

Sui, L., Mfopou, J.K., Geens, M., Sermon, K., and Bouwens, L. (2012). FGF signaling via MAPK is required early and improves Activin A-induced definitive endoderm formation from human embryonic stem cells. Biochem. Biophys. Res. Commun. 426, 380–385.

Vaninetti, N., Williams, L., Geldenhuys, L., Porter, G.A., Guernsey, D.L., and Casson, A.G. (2009). Regulation of CDX2 expression in esophageal adenocarcinoma. Mol. Carcinog. 48, 965–974.

Vorontsov, I.E., Fedorova, A.D., Yevshin, I.S., Sharipov, R.N., Kolpakov, F.A., Makeev, V.J., and Kulakovskiy, I. V. (2018). Genome-wide map of human and mouse transcription factor binding sites aggregated from ChIP-Seq data 06 Biological Sciences 0604 Genetics. BMC Res. Notes 11, 10–12.

Van Der Wal, E., Bergsma, A.J., Van Gestel, T.J.M., In ’t Groen, S.L.M., Zaehres, H., Araúzo-Bravo, M.J., Schöler, H.R., Van Der Ploeg, A.T., and Pim Pijnappel, W.W.M. (2017). GAA Deficiency in Pompe Disease Is Alleviated by Exon Inclusion in iPSC-Derived Skeletal Muscle Cells. Mol. Ther. Nucleic Acid 7, 101–115.

Wang, L., Xu, X., Cao, Y., Li, Z., Cheng, H., Zhu, G., Duan, F., Na, J., Han, J.-D.J., and Chen, Y.-G. (2017). Activin/Smad2-induced Histone H3 Lys-27 Trimethylation (H3K27me3) Reduction Is Crucial to Initiate Mesendoderm Differentiation of Human Embryonic Stem Cells. J. Biol. Chem. 292, 1339–1350.

Xie, W., Schultz, M.D., Lister, R., Hou, Z., Rajagopal, N., Ray, P., Whitaker, J.W., Tian, differentiation of human embryonic stem cells. Cell 153, 1134–1148.

Yanagihara, K., Liu, Y., Kanie, K., Takayama, K., Kokunugi, M., Hirata, M., Fukuda, T., Suga, M., Nikawa, H., Mizuguchi, H., et al. (2016). Prediction of Differentiation Tendency Toward Hepatocytes from Gene Expression in Undifferentiated Human Pluripotent Stem Cells. Stem Cells Dev. 25, 1884–1897.

Yang, S., Lin, G., Tan, Y.-Q., Deng, L.-Y., Yuan, D., and Lu, G.-X. (2010). Differences between karyotypically normal and abnormal human embryonic stem cells. Cell Prolif. 43, 195–206.

Zerbino, D.R., Achuthan, P., Akanni, W., Amode, M.R., Barrell, D., Bhai, J., Billis, K., Cummins, C., Gall, A., Girón, C.G., et al. (2018). Ensembl 2018. Nucleic Acids Res. 46, D754–D761.

