## Supplementary data for "Constitutional activation of BMP4 and WNT signalling in hESC results in impaired mesendoderm differentiation"

**Supplementary information**

**Table S1.** Top-25 upregulated and top-25 downregulated genes based on the Log_2_ fold-change between VUB03_S2 and the four control lines.

| ENSEMBL ID | Gene name | Log_2_FC | FDR |
| --- | --- | --- | --- |
| ENSG00000213401 | *MAGEA12* | 10.58 | 1.05E-17 |
| ENSG00000198930 | *CSAG1* | 9.14 | 2.16E-15 |
| ENSG00000221867 | *MAGEA3* | 8.42 | 1.46E-12 |
| ENSG00000144015 | *TRIM43* | 7.97 | 9.16E-11 |
| ENSG00000183305 | *MAGEA2B* | 7.90 | 0.000283051 |
| ENSG00000196589 | *MBD3L2B* | 7.47 | 1.28E-05 |
| ENSG00000197172 | *MAGEA6* | 7.27 | 1.18E-05 |
| ENSG00000268902 | *CSAG2* | 7.14 | 4.98E-11 |
| ENSG00000268916 | *CSAG3* | 7.14 | 4.98E-11 |
| ENSG00000180347 | *CCDC129* | 7.13 | 1.85E-12 |
| ENSG00000155622 | *XAGE2* | 6.75 | 6.95E-11 |
| ENSG00000203857 | *HSD3B1* | 6.68 | 0.000238888 |
| ENSG00000135346 | *CGA* | 6.58 | 1.55E-07 |
| ENSG00000268606 | *MAGEA2* | 6.19 | 2.09E-09 |
| ENSG00000182315 | *MBD3L3* | 5.94 | 1.22E-08 |
| ENSG00000116726 | *PRAMEF12* | 5.63 | 2.66E-08 |
| ENSG00000230522 | *MBD3L2* | 5.41 | 1.13E-06 |
| ENSG00000168930 | *TRIM49* | 5.36 | 2.59E-07 |
| ENSG00000203908 | *KHDC3L* | 5.29 | 2.70E-11 |
| ENSG00000169248 | *CXCL11* | 5.28 | 4.14E-08 |
| ENSG00000223417 | *TRIM49D1* | 5.27 | 1.02E-06 |
| ENSG00000184058 | *TBX1* | 5.23 | 6.01E-07 |
| ENSG00000144010 | *TRIM43B* | 4.90 | 2.24E-05 |
| ENSG00000228836 | *CT45A5* | 4.90 | 6.31E-05 |
| ENSG00000163993 | *S100P* | 4.71 | 1.75E-05 |
| ENSG00000203909 | *DPPA5* | 4.48 | 2.39E-12 |
| ENSG00000243137 | *PSG4* | 4.46 | 8.05E-05 |
| ENSG00000078401 | *EDN1* | 4.34 | 1.25E-06 |
| ENSG00000142182 | *DNMT3L* | 4.28 | 2.72E-05 |
| ENSG00000080007 | *DDX43* | 4.28 | 6.80E-09 |
| ENSG00000164744 | *SUN3* | 4.26 | 1.22E-08 |
| ENSG00000128253 | *RFPL2* | 4.25 | 1.07E-05 |
| ENSG00000089356 | *FXYD3* | 4.23 | 6.92E-08 |
| ENSG00000253117 | *OC90* | 4.22 | 4.09E-07 |
| ENSG00000164746 | *C7orf57* | 4.09 | 8.55E-07 |
| ENSG00000124721 | *DNAH8* | 3.99 | 3.20E-06 |
| ENSG00000068781 | *STON1-GTF2A1L* | 3.98 | 8.48E-06 |
| ENSG00000168453 | *HR* | 3.96 | 3.40E-05 |
| ENSG00000107954 | *NEURL1* | 3.96 | 1.43E-10 |
| ENSG00000186439 | *TRDN* | 3.93 | 3.12E-09 |
| ENSG00000185404 | *SP140L* | 3.88 | 1.45E-10 |
| ENSG00000188984 | *AADACL3* | 3.88 | 3.21E-07 |
| ENSG00000154645 | *CHODL* | 3.84 | 7.58E-11 |
| ENSG00000136167 | *LCP1* | 3.82 | 1.05E-06 |
| ENSG00000189134 | *NKAPL* | 3.81 | 5.74E-08 |
| ENSG00000113196 | *HAND1* | 3.76 | 1.71E-05 |
| ENSG00000179388 | *EGR3* | 3.75 | 1.76E-06 |
| ENSG00000182070 | *OR52A1* | 3.74 | 2.17E-07 |
| ENSG00000176945 | *MUC20* | 3.72 | 9.09E-06 |
| ENSG00000081853 | *PCDHGA2* | 3.66 | 7.61E-07 |
| ENSG00000049769 | *PPP1R3F* | -4.30 | 5.88E-11 |
| ENSG00000154864 | *PIEZO2* | -4.33 | 4.58E-05 |
| ENSG00000163661 | *PTX3* | -4.36 | 7.07E-05 |
| ENSG00000108405 | *P2RX1* | -4.43 | 0.005578534 |
| ENSG00000164708 | *PGAM2* | -4.44 | 1.07E-05 |
| ENSG00000128610 | *FEZF1* | -4.52 | 7.94E-05 |
| ENSG00000184227 | *ACOT1* | -4.53 | 0.024786873 |
| ENSG00000204071 | *TCEAL6* | -4.54 | 0.000291348 |
| ENSG00000054277 | *OPN3* | -4.55 | 9.48E-10 |
| ENSG00000172572 | *PDE3A* | -4.79 | 2.15E-05 |
| ENSG00000198822 | *GRM3* | -4.84 | 1.22E-08 |
| ENSG00000054803 | *CBLN4* | -4.85 | 0.073639428 |
| ENSG00000144057 | *ST6GAL2* | -4.86 | 1.03E-06 |
| ENSG00000198502 | *HLA-DRB5* | -4.88 | 0.105931456 |
| ENSG00000139364 | *TMEM132B* | -5.05 | 5.50E-08 |
| ENSG00000071991 | *CDH19* | -5.07 | 0.02685586 |
| ENSG00000043355 | *ZIC2* | -5.19 | 6.89E-08 |
| ENSG00000134463 | *ECHDC3* | -5.25 | 2.05E-08 |
| ENSG00000185985 | *SLITRK2* | -5.37 | 0.021596371 |
| ENSG00000229676 | *ZNF492* | -5.39 | 0.001028709 |
| ENSG00000173376 | *NDNF* | -5.76 | 0.000504175 |
| ENSG00000198739 | *LRRTM3* | -5.89 | 2.79E-05 |
| ENSG00000139800 | *ZIC5* | -5.93 | 1.41E-10 |
| ENSG00000177932 | *ZNF354C* | -6.09 | 4.73E-05 |
| ENSG00000092607 | *TBX15* | -6.24 | 0.020284181 |
| ENSG00000196932 | *TMEM26* | -6.26 | 3.00E-05 |
| ENSG00000166033 | *HTRA1* | -6.36 | 1.91E-11 |
| ENSG00000198028 | *ZNF560* | -6.39 | 0.002239187 |
| ENSG00000196109 | *ZNF676* | -6.47 | 0.005128662 |
| ENSG00000198105 | *ZNF248* | -6.60 | 0.043300948 |
| ENSG00000196767 | *POU3F4* | -6.71 | 3.76E-06 |
| ENSG00000140015 | *KCNH5* | -7.01 | 0.000126287 |
| ENSG00000212719 | *C17orf51* | -7.05 | 6.32E-08 |
| ENSG00000213973 | *ZNF99* | -7.12 | 0.000656566 |
| ENSG00000106153 | *CHCHD2* | -7.58 | 1.69E-06 |
| ENSG00000197134 | *ZNF257* | -7.68 | 7.25E-05 |
| ENSG00000160321 | *ZNF208* | -7.90 | 0.000169049 |
| ENSG00000277150 | *F8A3* | -8.81 | 0.012328624 |
| ENSG00000197360 | *ZNF98* | -8.93 | 0.004990358 |
| ENSG00000196350 | *ZNF729* | -9.87 | 0.002488076 |

**Table S2.** DAVID enrichment analysis of the top-1000 deregulated genes with fold enrichment >4.

| TERMS REFERRING TO CELL DIFFERENTIATION AND MORPHOGENESIS | P-Value | Fold Enrichment |
| --- | --- | --- |
| GO:0021537~telencephalon development | 1.18E-04 | 8.23137255 |
| GO:0061036~positive regulation of cartilage development | 0.00695347 | 6.24702381 |
| GO:0060384~innervation | 0.01079889 | 5.55291005 |
| GO:0001958~endochondral ossification | 0.00148785 | 5.38205128 |
| GO:0032332~positive regulation of chondrocyte differentiation | 0.01314789 | 5.26065163 |
| GO:0031290~retinal ganglion cell axon guidance | 0.01314789 | 5.26065163 |
| GO:0051480~regulation of cytosolic calcium ion concentration | 8.51E-04 | 4.99761905 |
| GO:0001892~embryonic placenta development | 0.01579845 | 4.99761905 |
| GO:0002062~chondrocyte differentiation | 5.81E-04 | 4.61318681 |
| GO:0030501~positive regulation of bone mineralization | 0.00149308 | 4.5692517 |
| GO:0090103~cochlea morphogenesis | 0.02204632 | 4.54329004 |
| GO:0035987~endodermal cell differentiation | 0.00992813 | 4.44232804 |
| GO:0003007~heart morphogenesis | 0.00453796 | 4.37291667 |
| GO:0003151~outflow tract morphogenesis | 3.86E-04 | 4.34575569 |
| GO:0035924~cellular response to vascular endothelial growth factor stimulus | 0.02566039 | 4.34575569 |
| GO:0009880~embryonic pattern specification | 0.02566039 | 4.34575569 |
| GO:0003148~outflow tract septum morphogenesis | 0.02566039 | 4.34575569 |
| GO:0045669~positive regulation of osteoblast differentiation | 3.41E-05 | 4.33126984 |
| GO:0045599~negative regulation of fat cell differentiation | 9.76E-04 | 4.28367347 |
| GO:0035116~embryonic hindlimb morphogenesis | 0.01160037 | 4.28367347 |
| TERMS REFERRING TO INTRACELLULAR SIGNALLING |  |  |
| GO:0017147~Wnt-protein binding | 9.25E-05 | 5.91900421 |
| GO:0005520~insulin-like growth factor binding | 0.01009102 | 5.66324477 |
| GO:0071837~HMG box domain binding | 0.01009102 | 5.66324477 |
| GO:0034199~activation of protein kinase A activity | 0.01079889 | 5.55291005 |
| GO:0071773~cellular response to BMP stimulus | 5.62E-04 | 5.33079365 |
| GO:0043425~bHLH transcription factor binding | 0.00448741 | 5.31852552 |
| GO:0005160~transforming growth factor beta receptor binding | 1.61E-04 | 4.8542098 |
| GO:0042813~Wnt-activated receptor activity | 0.02066408 | 4.6335639 |
| GO:0046427~positive regulation of JAK-STAT cascade | 0.02204632 | 4.54329004 |
| GO:0030513~positive regulation of BMP signaling pathway | 0.00384673 | 4.51397849 |
| GO:0032733~positive regulation of interleukin-10 production | 0.02566039 | 4.34575569 |
| GO:0010862~positive regulation of pathway-restricted SMAD protein phosphorylation | 5.38E-04 | 4.16468254 |
| TERMS REFERRING TO CELLULAR PROCESSES |  |  |
| GO:1901381~positive regulation of potassium ion transmembrane transport | 0.00542687 | 6.66349206 |
| GO:0006171~cAMP biosynthetic process | 0.00873885 | 5.87955182 |
| GO:0051412~response to corticosterone | 0.01079889 | 5.55291005 |
| GO:0001954~positive regulation of cell-matrix adhesion | 0.00397993 | 5.45194805 |
| GO:0007162~negative regulation of cell adhesion | 6.53E-05 | 5.4028314 |
| GO:0008373~sialyltransferase activity | 0.01229547 | 5.36517925 |
| GO:0032200~telomere organization | 0.00183313 | 5.18271605 |
| GO:0016339~calcium-dependent cell-cell adhesion via plasma membrane cell adhesion molecules | 0.00223583 | 4.99761905 |
| GO:2000352~negative regulation of endothelial cell apoptotic process | 0.00223583 | 4.99761905 |
| GO:0071385~cellular response to glucocorticoid stimulus | 0.01579845 | 4.99761905 |
| GO:0042326~negative regulation of phosphorylation | 0.01579845 | 4.99761905 |
| GO:0097503~sialylation | 0.01579845 | 4.99761905 |
| GO:0032870~cellular response to hormone stimulus | 3.25E-04 | 4.44232804 |
| GO:0006335~DNA replication-dependent nucleosome assembly | 0.00453796 | 4.37291667 |
| GO:0090398~cellular senescence | 0.02566039 | 4.34575569 |
| GO:0045786~negative regulation of cell cycle | 0.00209603 | 4.32226512 |
| GO:0040007~growth | 0.01160037 | 4.28367347 |

**Table S3.** List of the genetic content of lines and sublines used in the study

| **Line** | **Karyotype** | **Breakpoints (size)** |
| --- | --- | --- |
| VUB01 | 46, XY |  |
| VUB02 | 46, XY |  |
| VUB03_S1 | 46, XX |  |
| VUB03_S2 | 46, XX, dup(20)(q11.21) | 20: 31300536 – 35335783 (4Mb) |
| VUB14 | 46, XX |  |
| VUB19 | 46, XY |  |

**Table S4**. Type of de novo variants retrieved by exome sequencing of VUB03_S2, with read depth ≥ 10 and an allelic depth proportion ≥ 30%.

| **Type of Variant** | **Fraction of variants** |
| --- | --- |
| Intron | 30.3% |
| Sequence feature | 20.1% |
| Utr 3 prime | 17.2% |
| Upstream | 7.74% |
| Splice site region | 5.78% |
| Downstream | 4.91% |
| Non-synonymous coding | 3.60% |
| Utr 5 prime | 2.29% |
| Synonymous coding | 1.96% |
| Non-coding exon | 1.74% |
| Intergenic | 0.98% |
| Splice site acceptor | 0.76% |
| Codon change plus codon insertion | 0.44% |
| Codon change plus codon deletion | 0.44% |
| Frame shift | 0.44% |
| Tf binding site | 0.22% |
| Codon insertion | 0.22% |
| Stop gained | 0.22% |
| Within coding gene | 0.22% |
| Splice site donor | 0.22% |
| Start gained | 0.11% |
| Within non-coding gene | 0.11% |

**Table S5.** *De novo* non-synonymous changes in VUB03_S2 with read depth ≥ 10 and an allelic depth proportion ≥ 30%

| Gene symbol | Amino acid change | Base change | Polymorphism | Allelic depth proportion in VUB03_S2 | Read depth |
| --- | --- | --- | --- | --- | --- |
| *ADAMTS15* | p.Pro815Ser | c.2443C>T | Non synonymous | 31% | 13 |
| *AMHR2* | p.Glu28Lys | c.82G>A | Non synonymous | 62% | 37 |
| *APCDD1L* | p.Trp500Leu | c.1499G>T | Non synonymous | 59% | 27 |
| *ASPN* | p.Asp50_Asp51del | c.150_155delTGATGA | Codon change plus codon deletion | 56% | 18 |
| *ATXN3* | p.Gln292_Gln305dup | c.915_916insCAGCAGCAGCAGCAGCAGCAGCAGCAGCAGCAGCAGCAGCAG | Codon insertion | 48% | 50 |
| *CBX7* | p.Glu8* | c.22G>T | Stop gained | 30% | 10 |
| *CCDC159* | p.Gln136Lys | c.406C>A | Non synonymous | 56% | 36 |
| *CES1* | p.Ser12Ala | c.34T>G | Non synonymous | 100% | 18 |
| *CNTNAP3B* | p.Arg1214Pro | c.3641G>C | Non synonymous | 33% | 12 |
| *CRIPAK* | p.His132Asp | c.394C>G | Non synonymous | 67% | 21 |
| *CRIPAK* | p.Pro142Arg | c.425C>G | Non synonymous | 48% | 25 |
| *CRIPAK* | p.Pro173fs | c.517_518insGACGTGGAGTGCCCGCCTGCTCACACGTGCCCATGTGGAGTGCCCGCCTGCTCACACGTGCC | Frame shift | 89% | 18 |
| *CRTAP* | p.Arg212Leu | c.635G>T | Non synonymous | 50% | 34 |
| *DCTN1* | p.Gln514Lys | c.1540C>A | Non synonymous | 52% | 33 |
| *DSPP* | p.Ser1138_Ser1139dup | c.3416_3417insTGATAG | Codon change plus codon insertion | 100% | 40 |
| *DSPP* | p.Asn1140delinsSerAspSerSerAspSerSerAsp | c.3418_3419insGCGATAGCAGTGACAGCAGCG | Codon change plus codon insertion | 100% | 41 |
| *DSPP* | p.Glu1149_Ser1154dup | c.3461_3462insCGATAGCAGCGACAGCAG | Codon change plus codon insertion | 52% | 44 |
| *DSPP* | p.Ser1165_Asp1167del | c.3492_3500delTAGCAGCGA | Codon change plus codon deletion | 48% | 64 |
| *EPB41L2* | p.Val498Phe | c.1492G>T | Non synonymous | 34% | 29 |
| *FADS6* | p.Glu13Asp | c.17_18insGATGGAACCTACGGAGCCCATGGAACCTACGGAGCCCATGGAACCTACGGAGCCCATGGAACCTACGGAGCC | Codon change plus codon insertion | 100% | 22 |
| *FBRSL1* | p.Pro294Leu | c.881C>T | Non synonymous | 60% | 15 |
| *JAK3* | p.Ala797Glu | c.2390C>A | Non synonymous | 58% | 33 |
| *KRTAP5-5* | p.Gly44_Ala53del | c.129_158delAGGCTGTGGGGGCTGTGGCTCCGGCTGTGC | Codon change plus codon deletion | 76% | 75 |
| *LPIN1* | p.Glu13Asp | c.39G>C | Non synonymous | 100% | 10 |
| *LRP1B* | p.Leu1392Phe | c.4174C>T | Non synonymous | 35% | 31 |
| *MAP3K4* | p.Ala1199del | c.3596_3598delCTG | Codon change plus codon deletion | 74% | 38 |
| *MICAL1* | p.Ala12Thr | c.34G>A | Non synonymous | 50% | 18 |
| *MUC12* | p.Arg2634His | c.7901G>A | Non synonymous | 31% | 16 |
| *MUC2* | p.Thr1527Pro | c.4579A>C | Non synonymous | 65% | 182 |
| *MUC3A* | p.His15Pro | c.44A>C | Non synonymous | 30% | 98 |
| *MUC3A* | p.Ser19Ala | c.55T>G | Non synonymous | 31% | 105 |
| *MUC3A* | p.Ala22Gly | c.65C>G | Non synonymous | 33% | 116 |
| *MUC6* | p.Pro1963Thr | c.5887C>A | Non synonymous | 59% | 128 |
| *MUC6* | p.Arg1948Ser | c.5844A>C | Non synonymous | 37% | 124 |
| *MUC6* | p.Arg1948Gly | c.5842A>G | Non synonymous | 43% | 123 |
| *MUC6* | p.His1942Pro | c.5825A>C | Non synonymous | 50% | 118 |
| *NLRP6* | p.Pro244Ala | c.730C>G | Non synonymous | 45% | 11 |
| *NPIPB6* | p.Ala306dup | c.918_919insGTC | Codon insertion | 47% | 32 |
| *NPIPB6* | p.Ala306fs | c.916_917delGC | Frame shift | 43% | 30 |
| *PDCD6* | p.Ala12_Gly15del | c.34_45delGCCGGCCCTGGG | Codon deletion | 30% | 10 |
| *PLK1* | p.Asn430Lys | c.1290C>G | Non synonymous | 44% | 43 |
| *PPP1R9B* | p.Glu320* | c.958G>T | Stop gained | 40% | 10 |
| *RAB40B* | p.Gln264Pro | c.791A>C | Non synonymous | 30% | 50 |
| *RAD17* | p.Lys535Lys | c.1605G>A | Splice site region | 41% | 17 |
| *SLC22A18AS* | p.Asn223fs | c.666dupC | Frame shift | 36% | 107 |
| *SMIM13* | p.Leu35fs | c.104dupT | Frame shift | 30% | 30 |
| *TEX264* | p.Pro126Leu | c.377C>T | Non synonymous | 42% | 26 |
| *TLN2* | p.Tyr1739Phe | c.5216A>T | Non synonymous | 40% | 10 |
| *TNFRSF11A* | p.Met540Leu | c.1618A>C | Non synonymous | 30% | 20 |
| *WFS1* | p.Ser157Phe | c.470C>T | Non synonymous | 51% | 35 |
| *ZBTB7C* | p.Gln71His | c.213G>T | Non synonymous | 47% | 30 |

**Table S6.** Downstream genes and targets of *BMP4* and *CTNNB1*, containing a *de novo* variant in VUB03_S2

| Gene | Type of polymorphism | Reference | Alternative | Polymorphism effect |
| --- | --- | --- | --- | --- |
| *SFN* | DEL | GGTGTGT | GGT | UTR 3 prime |
| *GLI2* | INS | T | TACACAC | UTR 3 prime |
| *DPP4* | SNP | C | T | Sequence feature |
| *TLR9* | SNP | G | A | UTR 5 prime |
| *PRKCD* | INS | C | CAA | UTR 3 prime |
| *WNT5A* | DEL | CTTT | C | UTR 3 prime |
| *ESR1* | DEL | CAA | C | UTR 3 prime |
| *EPO* | DEL | CT | C | Sequence feature |
| *TCF7L2* | DEL | CTGTGTGTGTG | C | Sequence feature |
| *WT1* | INS | G | GGAGAGAGA | Intron |
| *CTNND1* | DEL | CT | C | Splice site region |
| *IGF1R* | INS | G | GA | Sequence feature |
| *PLK1* | SNP | C | G | Non synonymous |

**Table S7.** List of probes, assays and primers used for qRT-PCR.

| **Gene** | **Taqman Assay / Sequence** |
| --- | --- |
| *GUSB* | Hs99999908_m1 |
| *PAX6* | Hs00240871_m1 |
| *SOX17* | Hs00751752_s1 |
| *FOXA2* | Hs00232764_m1 |
| *PAX7* | Hs00242962_m1 |
| *RUNX1* | Hs00231079_m1 |
| *WNT3A* | Hs00902257_m1 |
| *MIXL1* | Hs05060541_s1 |
| *EOMES* | Hs00172872_m1 |
| *GSC* | Hs00418279_m1 |
| *CD144* | Hs00418279_m1 |
| *PECAM1* | Hs01065279_m1 |
| *FLK1* | Hs00911700_m1 |
| *CD117* | Hs00174029_m1 |
| *CD34* | Hs00990732_m1 |
| *KRT7* | Hs00559840_m1 |
| *CDX2* | Hs01078080_m1 |
| *OCT3A* | Forward 5’-GGA-CAC-CTG-GCT-TCG-GAT-TT-3’  Reverse 5’-CAT-CAC-CTC-CAC-CAC-CTG-G-3’  Probe 6-FAM- GCC-TTC-TCG-CCC-CC-MGB |
| *NANOG* | Forward 5’-TGC-AAA-TGT-CTT-CTG-CTG-AGA-TG-3’  Reverse 5’-TCC-TGA-ATA-AGC-AGA-TCC-ATG-GA-3’  Probe 6-FAM- CAG-AGA-CTG-TCT-CTC-CTC-MGB |
| *UBC* | Forward 5’-CGC-AGC-CGG-GAT-TTG-3’  Reverse 5’-TCA-AGT-GAC-GAT-CAC-AGC-GA-3’  Probe 6-FAM- TCG-CAG-TTC-TTG-TTT-GTG-MGB |

**Table S8**. List of antibodies used for immunofluorescence

| **Primary antibodies** | **Species** | **Company** | **Cat#** |
| --- | --- | --- | --- |
| PAX6 | Mouse Monoclonal IgG | Abcam | ab78545 |
| OCT3A | Mouse Monoclonal IgG | Santa Cruz | sc-5279 |
| OCT3A | Rabbit Monoclonal IgG | Cell Signalling | C30A3 |
| SOX17 | Goat Polyclonal IgG | R&D Systems | AF1924 |
| T | Goat Polyclonal IgG | R&D Systems | AF2085 |
| HNF4a | Mouse Monoclonal IgG | Santa Cruz | sc-374229 |
| PAX7 | Mouse Monoclonal IgG | R&D Systems | MAB1675 |
| **Secondary antibodies/**  **Fluorochrome** | **Species** | **Company** | **Cat#** |
| Alexa Fluor 488 | Goat anti-Mouse (H+L) | Thermo Fisher Scientific | A11001 |
| Alexa Fluor 488 | Donkey anti-Goat IgG (H+L) | Thermo Fisher Scientific | A11055 |
| Alexa Fluor 488 | Goat anti-Rabbit (H+L) | Thermo Fisher Scientific | A11034 |
| Alexa Fluor 546 | Donkey anti-Rabbit (H+L) | Thermo Fisher Scientific | A10040 |
| Alexa Fluor 594 | Donkey anti-Mouse IgG (H+L) | Thermo Fisher Scientific | R37115 |

**Figure S1.** KRT7 expression upon definitive endoderm differentiation of VUB03_S1 and VUB03_S2

Immunostaining for SOX17 (green) and POU5F1 (magenta) and KRT7 (red). Scale bars represent 100μm.

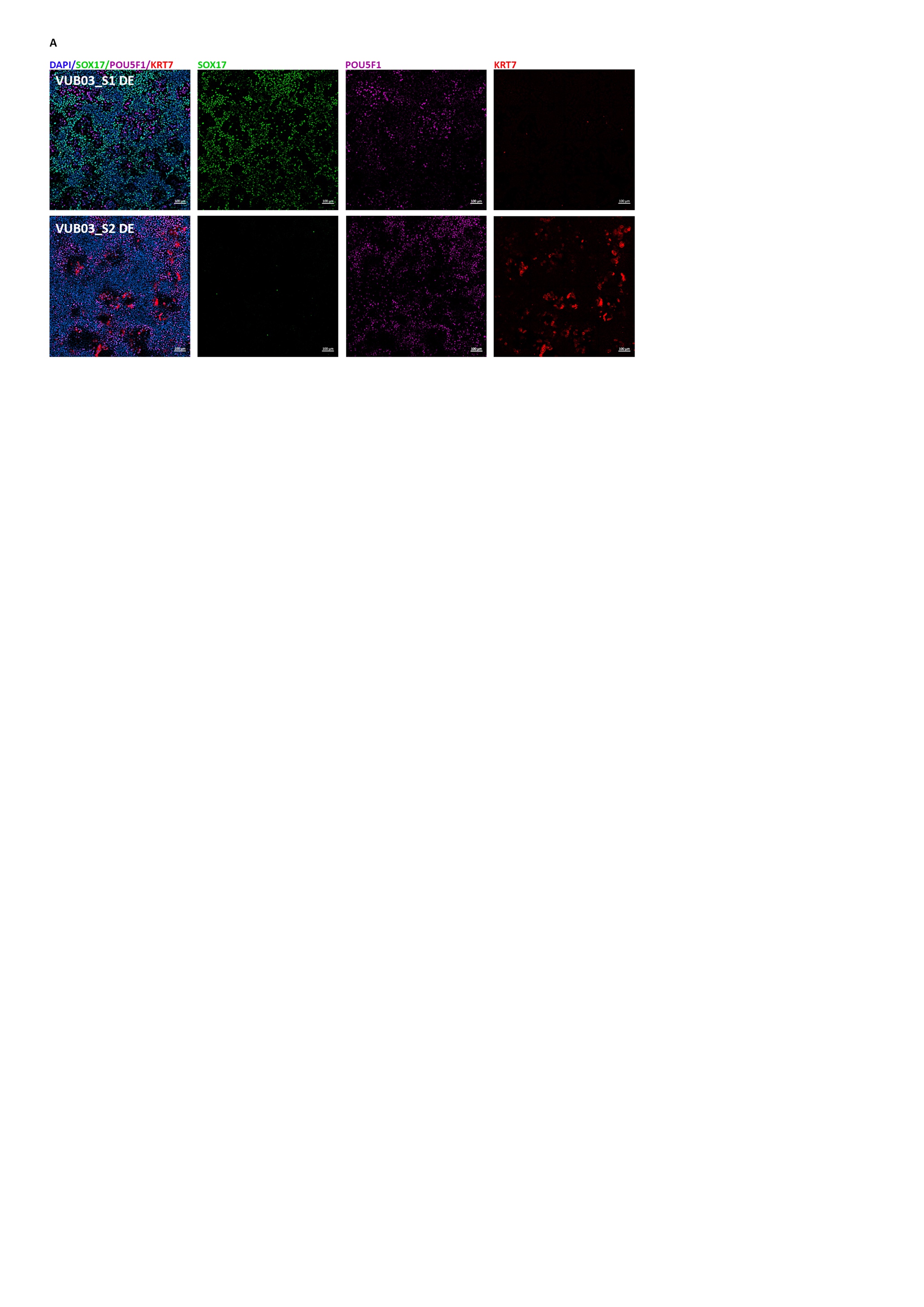

**Figure S2.** Impairing the definitive endoderm differentiation of VUB02 through WNT and BMP4 pathway activation. (A-B) mRNA expression of trophoblast markers *CDX2, KRT7* and definitive endoderm markers *SOX17* and *FOXA2* relative to VUB02 after definitive endoderm differentiation*.* (A) After use of BMP4 only. (B) After use of CHIR99021 only. (n=3) Data are shown as mean ±SEM, each dot represents an independent differentiation experiment. (C) Immunostaining for SOX17 (green) and POU5F1 (red) and (D) counts for SOX17 and POU5F1-positive cells in VUB02 after 3 days of definitive endoderm differentiation untreated and after BMP4 and CHIR treatment. Scale bars represent 2000μm and 100μm.

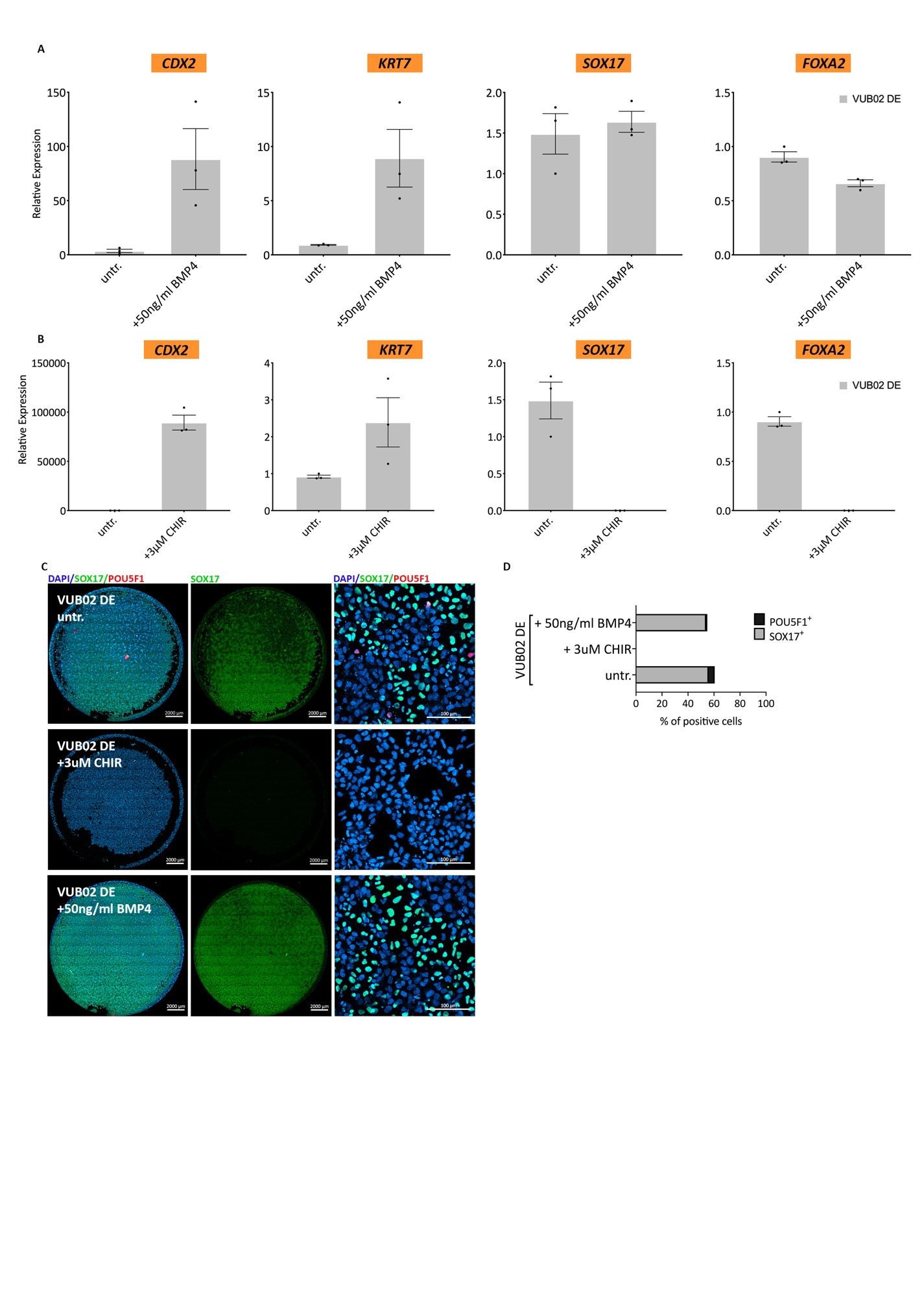

**Figure S3.** Impairing the definitive endoderm differentiation of VUB19 through WNT and BMP4 pathway activation. (A-B) mRNA expression of trophoblast markers *CDX2, KRT7* and definitive endoderm markers *SOX17* and *FOXA2* relative to VUB19 after definitive endoderm differentiation*.* (A) After use of BMP4 only. (B) After use of CHIR99021 only. (n=3) Data are shown as mean ±SEM, each dot represents an independent differentiation experiment. (C) Immunostaining for SOX17 (green) and POU5F1 (red) and (D) counts for SOX17 and POU5F1-positive cells in VUB19 after 3 days of definitive endoderm differentiation untreated and after BMP4 and CHIR treatment. Scale bars represent 2000μm and 100μm.

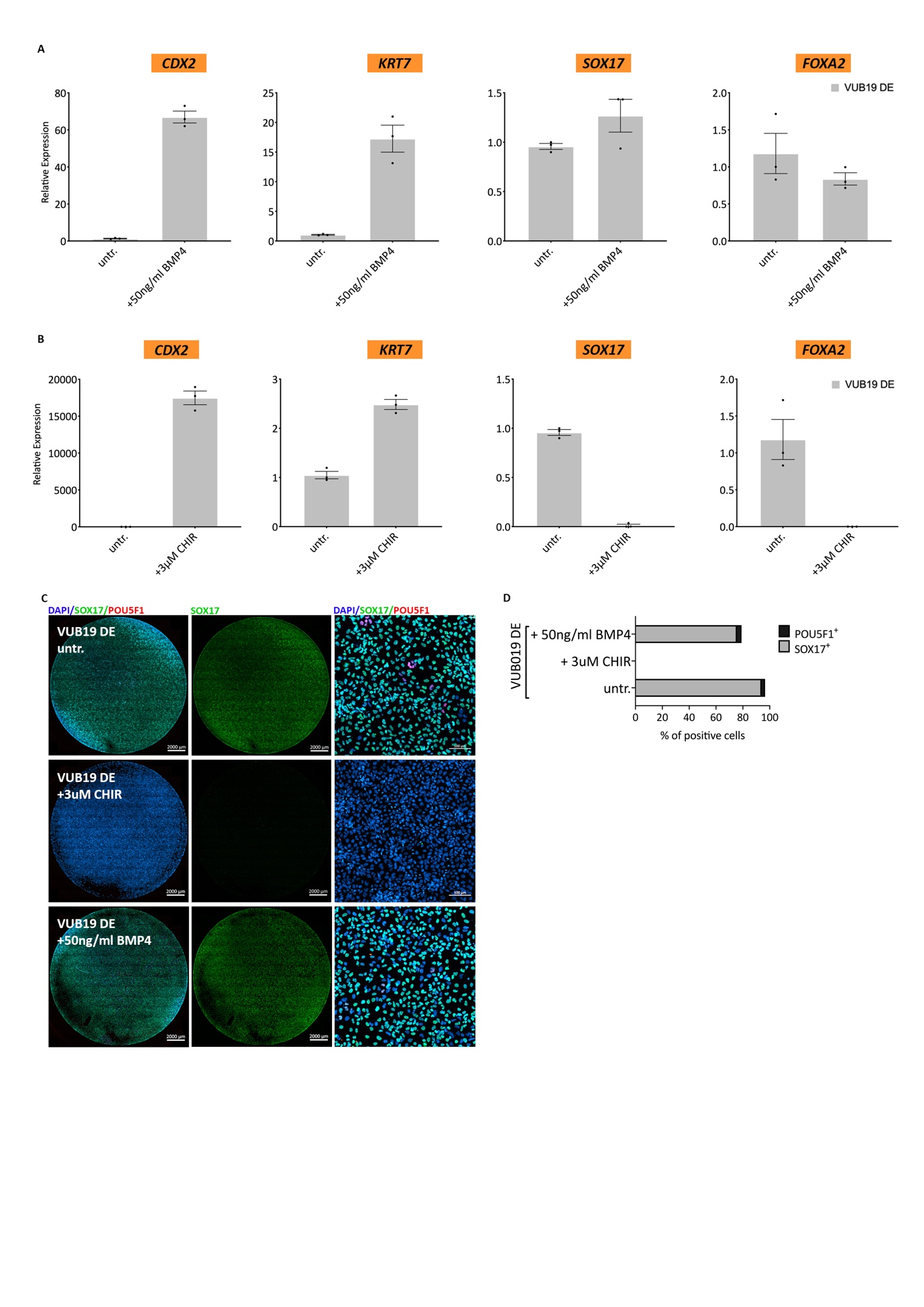
